# Phosphate selectivity in XPR1 is driven by direct multi-ion competition

**DOI:** 10.64898/2026.11.19.767951

**Authors:** Ciara Wallis, Xiaojie Wang, Yixiao Zhang, Ben Corry

## Abstract

Inorganic phosphate (Pi) is essential for numerous cellular processes, and its intracellular concentration must be tightly regulated. In mammals, phosphate efflux is mediated by the Xenotropic and Polytropic Retrovirus Receptor 1 (XPR1/SLC53A1), the only known mammalian Pi exporter. We and others have recently proposed that XPR1 functions like a Pi channel, with the protein preferentially conducting the H_2_PO_4_^-^ species of Pi over HPO_4_^2-^. While some studies suggest that XPR1 can conduct other anions, the molecular basis of XPR1 substrate selectivity remains unknown. Here, we use over 110 μs of molecular dynamics simulations, together with free-energy calculations, to investigate substrate selectivity in XPR1. We show that H_2_PO_4_^-^ (Pi^1-^) is preferentially conducted over HPO_4_^2-^ (Pi^1-^), and that Pi^1-^ progressively outcompetes Cl^-^ for occupancy of the transport pathway. Free-energy and ion-interaction analyses reveal that selectivity arises from two distinct mechanisms: favourable protein-Pi interactions and direct ion-ion interactions in the pore. This direct contact facilitates Pi^1-^ displacement via the previously proposed knock–kiss– kick mechanism, whereas entry of Cl^-^ is restricted due to the preference for water-mediated interactions. Simulations with additional anions show that XPR1 remains permissive to several monovalent species, including nitrate, bicarbonate and methanesulfonate, while poorly conducting the divalent anion sulfate. These findings suggest that XPR1 does not achieve phosphate selectivity through absolute exclusion of competing anions. Instead, preferential Pi transport emerges from substrate-specific interactions and competition between multiple ions within a relatively permissive permeation pathway, providing a molecular basis for substrate discrimination in XPR1.

## Introduction

Inorganic phosphate (Pi) is an important molecule in numerous biological processes,^1,2^ and regulating Pi levels within the cell is essential to prevent toxicity and metabolic dysfunction.^3^ Proteins from the SLC20 and SLC34 families are the major transporters responsible for Pi import into mammalian cells.^4–6^ In contrast, SLC53A1 (the Xenotropic and Polytropic Retrovirus Receptor 1; XPR1) is the only known mammalian protein that mediates Pi export.^7–9^ XPR1 is broadly expressed across various cell types and organs, and mutations have been linked to conditions such as ovarian clear cell carcinoma and other cancers, primary familial brain calcification, hypophosphataemia, and renal Fanconi syndrome.^10–17^

Structures of XPR1 reveal that the protein contains three key domains: the N-terminal SYG1/PHO/XPR1 (SPX) domain, the core transmembrane domain, and the C-terminal ERD1/XPR1/SYG1 (EXS) domain.^18–26^ The SPX domain functions as a sensor for inositol polyphosphates.^27^ In particular, InsP_8_ can bind to XPR1’s SPX domain, triggering conformational changes that enable Pi efflux to reduce intracellular Pi levels.^28–32^ We and others have suggested that human XPR1 and closely related homologues behave like channels, with Pi transport occurring through the core transmembrane and EXS domains.^18–26,33–35^ Our previous simulations of XPR1 indicated that dihydrogen phosphate (H_2_PO_4_^-^; Pi^1-^), not hydrogen phosphate (HPO_4_^2-^; Pi^2-^), was XPR1’s preferred substrate, and that substrate conduction occurred via a knock-kiss-kick mechanism in which Pi molecules transiently interact with charged residues lining the pore before the next incoming Pi displaces them.

The preference for Pi^1-^ over Pi^2-^ raises the question of how XPR1 discriminates between competing anions. Recent electrophysiological studies by Lu et al. (2024) and Wu et al. (2025) indicate that XPR1 can conduct other monovalent anions, including nitrate and methanesulfonate, but poorly conducts the divalent anion sulfate.^23,36^ However, the studies appear to differ in their observations of Cl^-^ conductance. Lu et al. (2024) found that XPR1 conducts Cl^-^ in the absence of Pi, whereas Wu et al. (2025) found that increasing extracellular Cl^-^ concentration did not alter XPR1 currents in the presence of Pi.^23,36^

To attempt to reconcile these differences and further examine XPR1 selectivity, we performed over 110 µs of molecular dynamics simulations of XPR1 in the presence and absence of different substrates. We show that Pi^1-^ outcompetes Cl^-^ for transport in our simulations, with few Cl^-^ permeation events occurring once Pi^1-^ occupies the pore. Further analysis of ion interactions and free energy calculations reveals two distinct mechanisms driving Pi selectivity. First, a specific binding site preferentially binds Pi due to highly favourable protein-ion interactions. Second, the unique ability of Pi, but not Cl^-^, to interact directly with other permeating ions allows it to efficiently displace resident ions from the pore. Simulations with competing anions further suggest that XPR1 preferentially transports phosphate while retaining the ability to conduct a range of other monovalent anions, whereas the divalent anion sulfate is not conducted.

Together, these findings suggest that the previously proposed knock-kiss-kick mechanism contributes to XPR1 substrate selectivity, explaining how the transporter preferentially conducts Pi while remaining permissive to other anions.

## Methods

Monomeric simulation systems of the intracellular-open/extracellular-open (iO-eO; PDB 9JXM) cryo-EM structure of XPR1 had been previously prepared^22^ for molecular dynamics (MD) simulations using the CHARMM-GUI Membrane Builder server.^37,38^ In these systems, only the transmembrane core and EXS domains (residues 230 to 625) were included to reduce system size. N- and C-terminal residues were acetylated and methylamidated, respectively. A disulfide bond was included between C415 and C440. The structures were inserted into a model membrane composed of pure 1,2-dioleoyl-sn-glycero-3-phosphocholine (DOPC) lipids. The position and orientation of XPR1 within the membrane was determined using the Predictions of Proteins in Membranes (PPM v2.0) server^39^, with the main axis of the protein aligned along the simulation box’s Z-axis. Systems were initially solvated with 0.15 M NaCl solution. The structural lipid in each monomer between TM2, TM4, and TM5 was added and modelled as DOPC in coot 0.9.7^40^ and further refined in real space using Phenix 1.21.1.^41^ The overall system dimensions were approximately 10 × 10 × 12 nm^3^, and consisted of ∼100,000 atoms, including ∼200 DOPC lipids, ∼60 Na^+^ ions, ∼70 Cl^-^ ions, and ∼21,500 water molecules.

The protonation states of all aspartate, glutamate, and histidine residues were set to the states we previously predicted using continuous constant pH MD simulations.^22^

As described previously, all simulations were performed using the GPU-accelerated version of GROMACS (v.2023)^42^ with the CHARMM36m forcefield^43^ with WYF parameters for cation-pi interactions^44^, and the TIP3P water model. Steepest descent energy minimisation was performed with a tolerance of 1,000 kJ mol^−1^ nm^−1^. During minimisation, all protein backbone and sidechain atoms were position-restrained with force constants of 4,000 kJ mol^−1^ nm^−1^ and 2,000 kJ mol^−1^ nm^−1^, respectively. Position restraints of 1,000 kJ mol^−1^ nm^−1^ were also placed on the phosphorous atoms of DOPC headgroups. After minimising, systems were equilibrated over six steps for a total of 60 ns, where position restraints were gradually released on all atoms, with only the Cα atoms of residues 618-625 remaining restrained with a force constant of 50 kJ mol^−1^ nm^−1^ to maintain the orientation of the C-Loop. The LINCS algorithm was used to constrain covalent bonds to hydrogen atoms, allowing a 2 fs timestep to be used.^45^

Electrostatic interactions within a cut-off of 1.2 nm were calculated using the Particle-Mesh Ewald algorithm^46^, using the Verlet grid cut-off scheme with an update frequency of every 20 steps. Long-range van der Waals interactions were calculated within a cut-off of 1.2 nm, with a smooth switching function beginning from 1.0 nm. All simulations were performed under periodic boundary conditions. In the equilibration stages, the Berendsen thermostat was used to maintain the simulation temperature at 310.15 K, using a time constant of 1 ps.^47^ Three thermally coupled groups consisting of 1) the protein, 2) the membrane and 3) the water, Na^+^, Cl^-^, and Pi ions were used to improve the accuracy of the simulated temperature. The Berendsen barostat^47^ was used to maintain simulation pressure at 1 bar, using a time constant of 5 ps with a compressibility of 4.5e-5 bar^-1^. During production runs, the v-rescale thermostat^48^ and Parrinello-Rahman barostat^49^ were used to semi-isotropically maintain the temperature and pressure at 310.15 K and 1.0 bar, respectively.

The structures of dihydrogen phosphate (H_2_PO_4_^-^; CID 1003), hydrogen phosphate (HPO_4_^2-^; CID 3681305), methanesulfonate (CH_3_SO_3_^-^; CID 85257), bicarbonate (HCO_3_^-^; CID 769), nitrate (NO_3_^-^; CID 943), and sulfate (SO_4_^2-^; CID 1118) were sourced from PubChem and were geometry optimised via quantum mechanics calculations using the Automated Topology Builder Webserver.^50^ Parameters for these substrates were obtained using the CHARMM General Force Field (CGenFF).^51^ To improve the chances of observing substrate permeation, an electric field was applied during production simulations along the Z-axis of the system with strengths of -0.035 V nm^−1^ to create a potential difference across the simulation box of approximately -400 mV. Systems were simulated with five replicates for 2 μs per replicate, with the exception of the simulations that contained only Pi^1-^ from Wang et al. (2025).^22^ Different combinations of anions (dihydrogen phosphate, hydrogen phosphate, methanesulfonate, bicarbonate, and nitrate) were added to the simulation systems. The number of anion species added to each system is included in **Table 1**. Systems were charge neutralised using GMX GENION.

**Table 1.** Substrate selectivity simulation systems used in this study. All simulations were performed in the presence of an electric field, mimicking a transmembrane potential of -400 mV. Cl^-^ ions were present in all systems. Simulation systems from Wang et al. (2025) are italicised and shown in light grey.^22^

| XPR1 Conformation | Pi Species | Other Anion | Replicates | Duration Per Replicate | Total Duration |
|---|---|---|---|---|---|
| iO-eO | N/A | N/A | 5 | 2 µs | 10 µs |
| *iO-eO* | *H<sub>2</sub>PO<sub>4</sub><sup>-</sup> (25)* | *N/A* | *4* | *2 µs* | *8 µs* |
| *iO-eO* | *H<sub>2</sub>PO<sub>4</sub><sup>-</sup> (12), HPO<sub>4</sub><sup>2-</sup> (12)* | *N/A* | *5* | *2 µs* | *10 µs* |
| iO-eO | HPO<sub>4</sub><sup>2-</sup> (25) | N/A | 5 | 2 µs | 10 µs |
| iO-eO | N/A | CH<sub>3</sub>SO<sub>3</sub><sup>-</sup> (25) | 5 | 2 µs | 10 µs |
| iO-eO | N/A | HCO<sub>3</sub><sup>-</sup> (25) | 5 | 2 µs | 10 µs |
| iO-eO | N/A | NO<sub>3</sub><sup>-</sup> (25) | 5 | 2 µs | 10 µs |
| iO-eO | N/A | SO<sub>4</sub><sup>2-</sup> (25) | 5 | 2 µs | 10 µs |
| iO-eO | H<sub>2</sub>PO<sub>4</sub><sup>-</sup> (12) | CH<sub>3</sub>SO<sub>3</sub><sup>-</sup> (12) | 5 | 2 µs | 10 µs |
| iO-eO | H<sub>2</sub>PO<sub>4</sub><sup>-</sup> (12) | HCO<sub>3</sub><sup>-</sup> (12) | 5 | 2 µs | 10 µs |
| iO-eO | H<sub>2</sub>PO<sub>4</sub><sup>-</sup> (12) | NO<sub>3</sub><sup>-</sup> (12) | 5 | 2 µs | 10 µs |

Conductance was calculated from intervals of each simulation replicate using the following equation, where N is the total number of permeation events observed in the simulation interval, q is the charge carried by each permeating ion (−1e for monovalent anions and −2e for divalent anions), t is the simulation time in seconds, and V is the transmembrane potential in volts.

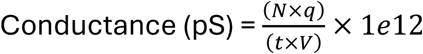

As transitions from the conductive iO-eO state towards the less conductive iO-eC conformation were observed within the 2 μs simulation timescale, conductance calculated across the complete trajectory would include periods in which the protein had transitioned away from its starting conformation. To estimate the conductance associated with the iO-eO state, each trajectory was divided into seven overlapping 500 ns intervals: 0–500 ns, 250– 750 ns, 500–1000 ns, 750–1250 ns, 1000–1500 ns, 1250–1750 ns, and 1500–2000 ns.

Conductance was calculated separately for each interval, and the maximum 500 ns conductance was used as an estimate of the conductive capacity of the iO-eO state for each independent replicate. This procedure was applied consistently across all simulation systems to allow comparisons between different substrates. The replicate-level estimates were then averaged for each simulation system.

To further explore the molecular basis of selectivity for Pi over Cl^-^, free energy perturbation (FEP) simulations were performed in GROMACS (v.2023)^42,52^ for the Pi-0 and Pi-1 sites shown in **Fig. 2C**. For each site, a representative structure was selected from the iO-eO simulations in which Pi occupied that site. A Cl^-^ ion was then placed at the position of the phosphorus atom, and a hybrid Cl^-^–phosphate topology was generated in which Pi and Cl^-^ represented the two alchemical end states. In the hybrid topology, the Cl^-^ atom was present in one end state, while the atoms of H_2_PO_4_^-^ were present in the alternate end state as dummy atoms. Bonded interactions for the phosphate group were retained throughout the transformation to preserve its internal geometry. Both forward (Cl^-^→Pi) and reverse (Pi→Cl^-^) transformations were performed for each binding site. Free-energy perturbation systems are shown in **Table 2**.

**Table 2.** Summary of free energy perturbation simulations performed in this study.

| **Site** | **Environment** | **Transformation** | **Replicates** | **λ windows** | **Production per λ** | **Total production** |
|---|---|---|---|---|---|---|
| Pi-1 | Protein | Cl<sup>−</sup> → Pi | 3 | 19 | 10 ns | 570 ns |
| Pi-1 | Protein | Pi → Cl<sup>−</sup> | 3 | 19 | 10 ns | 570 ns |
| Pi-0 | Protein | Cl<sup>−</sup> → Pi | 3 | 19 | 10 ns | 570 ns |
| Pi-0 | Protein | Pi → Cl<sup>−</sup> | 3 | 19 | 10 ns | 570 ns |
| Pi-0 | Protein | Cl<sup>−</sup> → Pi (with Pi at Pi-1) | 3 | 19 | 10 ns | 570 ns |
| Pi-0 | Protein | Pi → Cl<sup>−</sup> (with Pi at Pi-1) | 3 | 19 | 10 ns | 570 ns |
| Bulk | Water box | Cl<sup>−</sup> → Pi | 3 | 19 | 10 ns | 570 ns |
| Bulk | Water box | Pi → Cl<sup>−</sup> | 3 | 19 | 10 ns | 570 ns |

All additional Pi molecules were removed from the protein system prior to the FEP setup and were replaced with Cl^-^ ions, unless specified otherwise. To prevent bulk ions entering the transport pathway during the simulations, an additional dummy exclusion atom was positioned 4 Å from the hybrid ligand and assigned repulsive Lennard–Jones interactions with bulk Cl^-^ and Na⁺ ions (σ = 1.2 nm, ε = 5.0 kJ mol^-1^). To prevent the Cl^-^ atom of the hybrid ligand from also being repelled, it was assigned a separate atom type with identical nonbonded parameters to bulk Cl^-^. To maintain the position of the hybrid ligand within the binding site, harmonic position restraints of 500 kJ mol^-1^ nm^-1^ were applied to the Cl^-^, phosphorus, and dummy atoms in the x, y and z directions.

Equivalent forward and reverse transformations were also performed in bulk solvent using a 30 × 30 × 30 Å³ box of TIP3P water molecules, containing the hybrid ligand and a single Na⁺ counterion. A 30 × 30 × 30 Å³ TIP3P water box has been previously used for FEP calculations in bulk by Thomas, Jayatilaka, and Corry.^53^ The same positional restraints described above for protein transformations were also applied to the hybrid ligand during the bulk solvent transformations. Protein and bulk-solvent transformations were performed across 19 λ windows (0.00, 0.01, 0.02, 0.05, 0.10, 0.15, 0.20, 0.30, 0.40, 0.50, 0.60, 0.70, 0.80, 0.90, 0.925, 0.95, 0.98, 0.99 and 1.00). Window spacing was chosen to ensure sufficient overlap between neighbouring λ windows. Protein systems were energy minimised and equilibrated as described above for the standard molecular dynamics simulations. Bulk solvent systems were energy minimised, followed by 1 ns NVT and 2.5 ns NPT equilibration using the Berendsen thermostat and barostat. Production simulations were then performed for 10 ns at each λ window, with three independent replicates for both the protein and bulk solvent systems.

Free energy differences between adjacent λ windows were estimated using the Bennett Acceptance Ratio (BAR) method as implemented in GROMACS.^42,54^ Relative binding free energies were calculated as ΔΔG = ΔG protein – ΔG bulk, where ΔG protein and ΔG bulk are the free energy changes associated with the alchemical transformation in the protein and bulk-solvent systems, respectively (**Table S1**). Convergence was assessed by comparing independent replicates and agreement between the forward (Cl^-^ to Pi) and reverse (Pi to Cl^-^) transformations (**Fig. S4**). Additional FEP simulations were performed where Pi^1-^ was restrained at the Pi-1 site and the hybrid Pi-Cl^-^ ligand was present at the Pi-0 site. The Pi^1-^ molecule was restrained at the Pi-1 site using harmonic position restraints of 500 kJ mol^-1^ nm^-2^, applied to the phosphorus atom in the x, y and z directions.

These simulations were performed in triplicate for both the Cl^-^ → Pi and Pi → Cl^-^ transformations, using the same protocol as the other FEP simulations.

Per-residue MM/GBSA analysis was performed using gmx_MMPBSA (v1.6.5).^55,56^ Protein-only, ligand-only, and protein–ligand complex trajectories were generated from each replicate of the forwards FEP simulations at the Pi-0 and Pi-1 sites from the λ = 0 (window 0, Cl^-^ endpoint) and λ = 1 (window 18, Pi^1-^ endpoint) windows using gmx trjconv. For the λ = 0 windows, only the Cl^-^ atom from the hybrid Pi-Cl^-^ molecule was included when stripping the trajectory with gmx trjconv. Likewise, for the λ = 1 windows, only the atoms corresponding to Pi^1-^ were included. To prevent the Cl^-^ atom from being recognised as solvent and removed during gmx_MMPBSA preparation, the atom was renamed to allow calculations to be performed. MM/GBSA calculations were run with the Generalised Born implicit solvent model (igb = 5) with an implicit salt concentration of 150 mM. Per-residue free energy decomposition was performed (idecomp = 1) for residues within 8 Å of the ligand. Analysis was performed on all frames from the 10 ns trajectories. Per-residue interaction energies for Pi^1-^ and Cl^-^ at the Pi-0 and Pi-1 sites are reported as the mean ± SEM of three independent simulations.

Analysis of Pi permeation events were performed using a script modified from Adamson et al.^57^ The number of permeation events from monomeric systems was doubled to approximate expected permeation events through an XPR1 dimer. Systems were visualized using the Visual Molecular Dynamics software.^58^ All other analysis and graphing were performed in Python using the MDAnalysis^59^, Pandas^60^, NumPy^61^, Matplotlib^62^, and Seaborn libraries^63^. Statistical analysis was performed using SciPy.^64^

## Results

In our previous simulations of XPR1, we showed that Pi^1-^ outcompeted Pi^2-^ when both species were present. However, whether Pi^2-^ could permeate XPR1 in the absence of Pi^1-^ remained unclear. To investigate this, we performed simulations containing Pi^2-^ alone and compared these with our previous simulations containing Pi^1-^ alone or both phosphate species. Notably, Pi^2-^ permeation events do not increase in the absence of Pi^1-^, further supporting Pi^1-^ as the preferred Pi species conducted by XPR1 (**Fig. 1A**).

**Figure 1.**
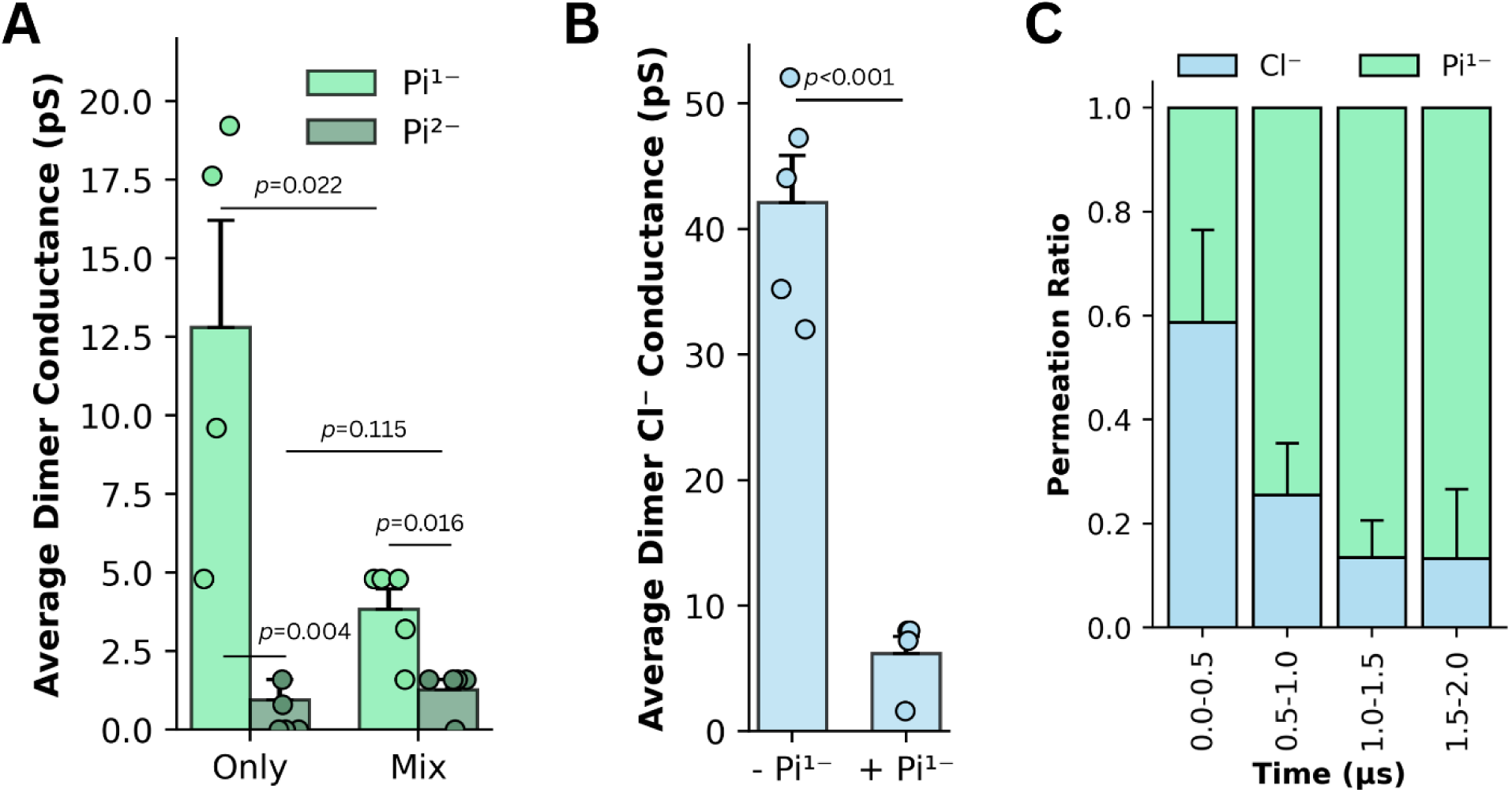
XPR1 preferentially conducts H_2_PO_4_^-^ over HPO_4_^2-^ and Cl^-^. **A)** Average dimer conductance (pS) for H_2_PO_4_^-^ (Pi^1-^) and HPO_4_^2-^ (Pi^2-^) from simulations containing only a single phosphate species (“Only”) or both phosphate species together (“Mix”). Statistical significance was assessed using unpaired t-tests for comparisons between the two “Only” systems and between “Only” and “Mix” systems, while paired t-tests were used to compare Pi^1-^ and Pi^2-^ conductance within the mixed simulations. **B)** Average Cl^-^ conductance from simulations performed in the absence (− Pi^1-^) and presence (+ Pi^1-^) of H_2_PO_4_^-^. Statistical significance was assessed using an unpaired t-test. For **A** and **B**, bars represent the mean conductance for each simulation system, with scatter points used to show the data from each replicate. Error bars represent SEM. The *p* values are shown above each comparison. **C)** Relative permeation ratios of H_2_PO_4_^-^ and Cl^-^ over successive time intervals from simulations containing both anion species. Error bars represent SEM.

Intriguingly, in the absence of either Pi species, a substantial number of Cl^-^ permeation events were observed. Cl^-^ permeation also occurred in simulations where Pi was present, but to a significantly lesser degree than when Cl^-^ was the only anion present (**Fig. 1B**).

When we examine the ratio of Pi^1-^ to Cl^-^ permeation events from different intervals of the simulation containing both anions, we observe that progressively fewer Cl^-^ permeations occur over the course of the simulation. These data suggest that Pi^1-^ outcompetes Cl^-^ for the transport pathway (**Fig. 1C, Movie S1**).

Our previously proposed knock-kiss-kick mechanism suggests that Pi molecules come in close proximity to each other to get displaced and conducted through the transport pathway.^22^ As Cl^-^ ions are unlikely to exhibit similar behaviour, we examined the spatial distributions and interactions of Pi^1-^ and Cl^-^ within the transport pathway to investigate the structural basis of Pi^1-^ selectivity.

We generated 2D density maps from simulations containing only Cl^-^ and from the final 1 μs of simulations containing Pi^1-^, by which point Cl^-^ had mostly been replaced by Pi^1-^ within the cavity. In the absence of Pi, Cl^-^ density was observed throughout the transport pathway, peaking around the Pi-1, Pi-2 and Pi-3 sites (**Fig. 2A**). However, when Pi^1-^ was present, Cl^-^ density largely disappeared (**Fig. 2B**), and Pi^1-^ occupied these positions instead. Additionally, Pi^1-^ was also found to occupy a fourth site immediately below the Pi-1 site on the intracellular side, which we call Pi-0, (**Fig. 2C**).

**Figure 2.**
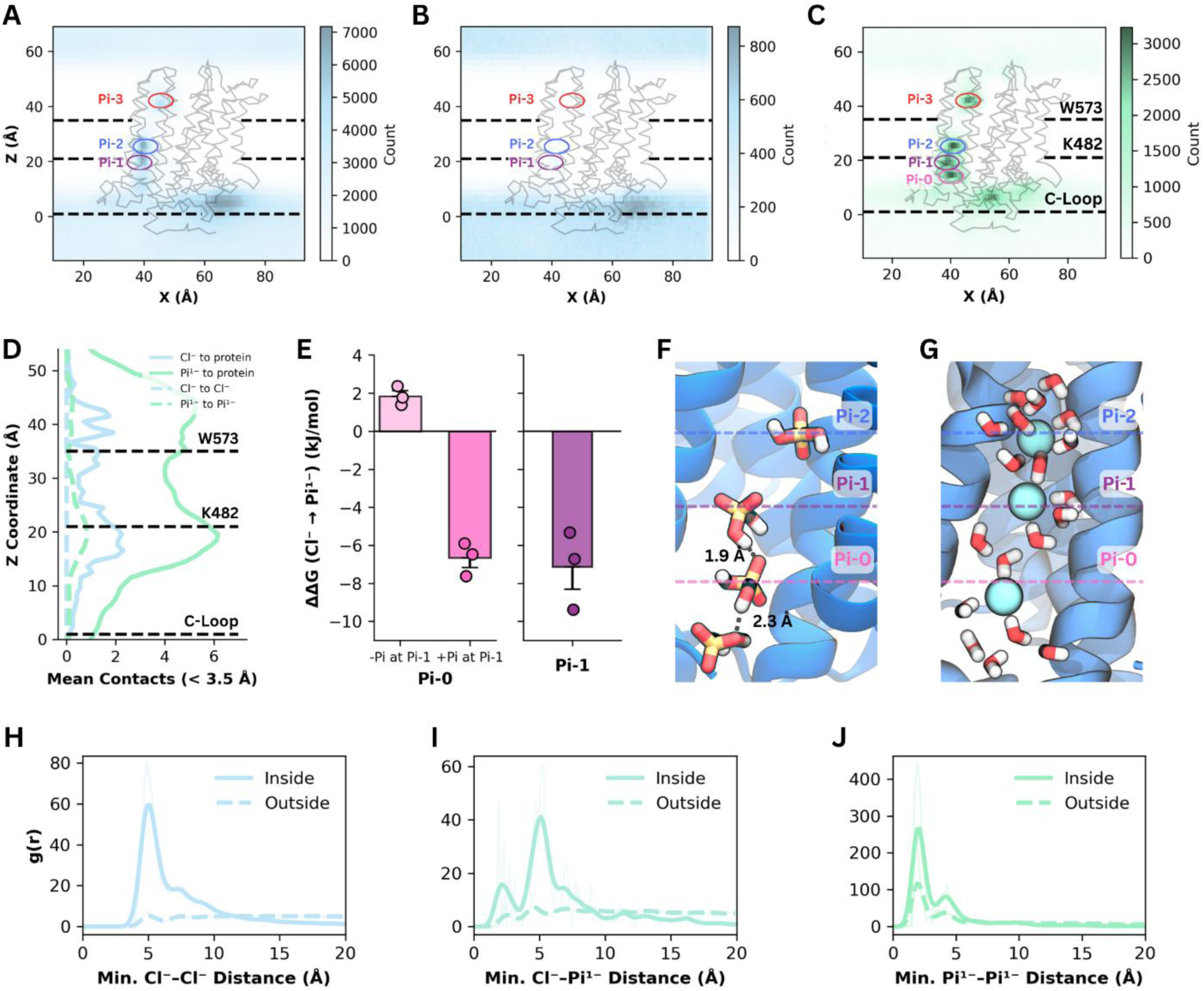
Phosphate outcompetes Cl^-^ via a multi-ion exclusion mechanism. **A–C)** Two-dimensional density maps showing the locations sampled by **A)** Cl^-^, in simulations where Pi^1-^ was absent, **B)** Cl^-^, from simulations where Pi^1-^ was present, and **C)** Pi^1-^ during simulations. The locations of the Pi-1, Pi-2, and Pi-3 sites are shown with purple, blue, and red circles, respectively. An additional Pi-0 site in **C** is shown with a pink circle. For **A-C**, data from all replicates was combined to make the density distributions. Dashed lines indicate the positions of the intracellular C-Loop, K482, and W573. **D)** Mean number of contacts (<3.5 Å) formed by Cl^-^ and Pi^1-^ with the side chains of pathway residues (solid lines) and with ions of the same species (dashed lines) as a function of position along the transport pathway. **E)** Relative binding free energies (ΔΔG, kJ/mol) for the Cl^-^ → Pi transformation at the Pi-0 and Pi-1 binding sites, determined using free energy perturbation (FEP) simulations. FEP simulations at Pi-0 were performed both in the absence and presence of Pi at the Pi-1 site. Bars represent the mean of three independent replicates, with individual replicate values overlaid. Error bars represent the standard error of the mean (SEM). **F-G)** Representative snapshot illustrating the formation of a chain of **F)** hydrogen-bonding Pi^1-^ molecules and **G)** Cl^-^ ions separated by water molecules within the transport pathway. Hydrogen bonds are shown as dashed black lines. Distances between adjacent phosphate ions are indicated. The locations of the Pi-0, Pi-1, and Pi-2 sites are shown as dashed pink, purple, and blue lines, respectively. **H–J)** Radial distribution functions showing the distributions of minimum **H)** Cl^-^–Cl^-^, **I)** Cl^-^–Pi^1-^, and **J)** Pi^1-^–Pi^1-^ distances for ions inside and outside the transport pathway. Traces for outside the pathway in **H-J** were multiplied by 5 to be more visible within the panel.

Quantification of the mean contacts formed with XPR1 and neighbouring ions (**Fig. 2D**) showed that Pi^1-^ generally formed more contacts with the protein than Cl^-^, assembling into hydrogen-bonded Pi^1-^ chains throughout the cavity (**Fig. 2F**). In contrast, Cl^-^ ions remained separated by intervening water molecules (**Fig. 2G**).

To explore the origin of this Pi selectivity, we performed free energy perturbation (FEP) calculations at the Pi-0 and Pi-1 sites (**Fig. 2E**). In the absence of other ions in the pathway, the calculated free energy difference between Cl^-^ and Pi^1-^ at the Pi-0 site was small (ΔΔG = +1.85 ± 0.26 kJ/mol), indicating little inherent discrimination. Conversely Pi^1-^ binding at the Pi-1 site was substantially more favourable than Cl^-^ (ΔΔG = −7.13 ± 1.17 kJ/mol). At both sites, Pi^1-^ formed more interactions with surrounding protein residues than Cl^-^ (**Fig. S5-8**). MM/GBSA analysis supported that Pi^1-^ formed more favourable interactions at Pi-1, primarily driven by stronger coordination with R570, R603, R604, and T525 (**Fig. S8A-B**).

To investigate whether occupancy of the Pi-1 site influences the adjacent Pi-0 site, we performed additional FEP simulations with Pi^1-^ restrained at Pi-1. Under these conditions, the relative preference at Pi-0 shifted dramatically in favour of Pi^1-^ (ΔΔG = −6.66 ± 0.88 kJ/mol). This cooperative effect explains why Cl^-^ occupancy at Pi-0 was not observed in the presence of Pi^1-^ (**Fig. 2B**), indicating that phosphate occupancy at Pi-1 actively drives the exclusion of Cl^-^ from the adjacent site.

Finally, we investigated how incoming ions enter the occupied pathway by calculating radial distribution functions for ion pairs inside the pore and in bulk solution. Within the transport pathway, the minimum Cl^-^ to Cl^-^ and Pi^1-^ to Cl^-^ profiles exhibit a major peak at approximately 5 Å, consistent with a water molecule bridging the two ions (**Fig. 2G-I**). For Pi^1-^ - Pi^1-^ pairs, however, the dominant peak shifted to 2 Å (**Fig. 2J**).

These findings demonstrate that once a Pi^1-^ occupies the Pi-1 site, a subsequent incoming Pi^1-^ can sit directly adjacent to it at Pi-0 through close favourable Pi-Pi interactions. These direct interactions facilitate the displacement of the bound Pi^1-^ via our previously proposed knock–kiss–kick mechanism. In contrast, Cl^-^ rarely occupies Pi-0 while Pi^1-^ resides at Pi-1 (**Fig. 2B-C**). Because Cl^-^ maintains an intervening water, its interactions are insufficient to promote regular displacement of Pi^1-^ from the Pi-1 site, effectively blocking Cl^-^ entry into the pathway once Pi^1-^ has taken up residence. Together, these results indicate that substrate selectivity depends on the ability of neighbouring Pi^1-^ ions to form direct interactions without intervening water molecules. This close proximity drives ion displacement via the proposed knock–kiss–kick mechanism, whereas Cl^-^ is less able to displace bound ions because it maintains an intervening shell of water molecules.

After investigating the molecular basis of XPR1 selectivity for Pi^1-^ over Pi^2-^ and Cl^-^, we examined whether XPR1 prefers Pi^1-^ over other divalent and monovalent anions. Previous electrophysiology studies by Lu et al. (2024) and Wu et al. (2025) have shown that XPR1 can conduct the monovalent anions methanesulfonate and nitrate, but poorly conducts sulfate.^23,36^ We performed MD simulations of XPR1 with methanesulfonate, sulfate, nitrate, and the additional monovalent anion bicarbonate (**Fig. 3A**, **Fig. S1A-E**). In the absence of Pi^1-^, the conductance values of methanesulfonate, bicarbonate and nitrate were not significantly different from those measured for Pi^1-^ in simulations. In contrast, sulfate showed very few permeation events, comparable to those observed for Pi^2-^, suggesting that XPR1 may be selective against divalent anions (**Fig. 3B**).

**Figure 3.**
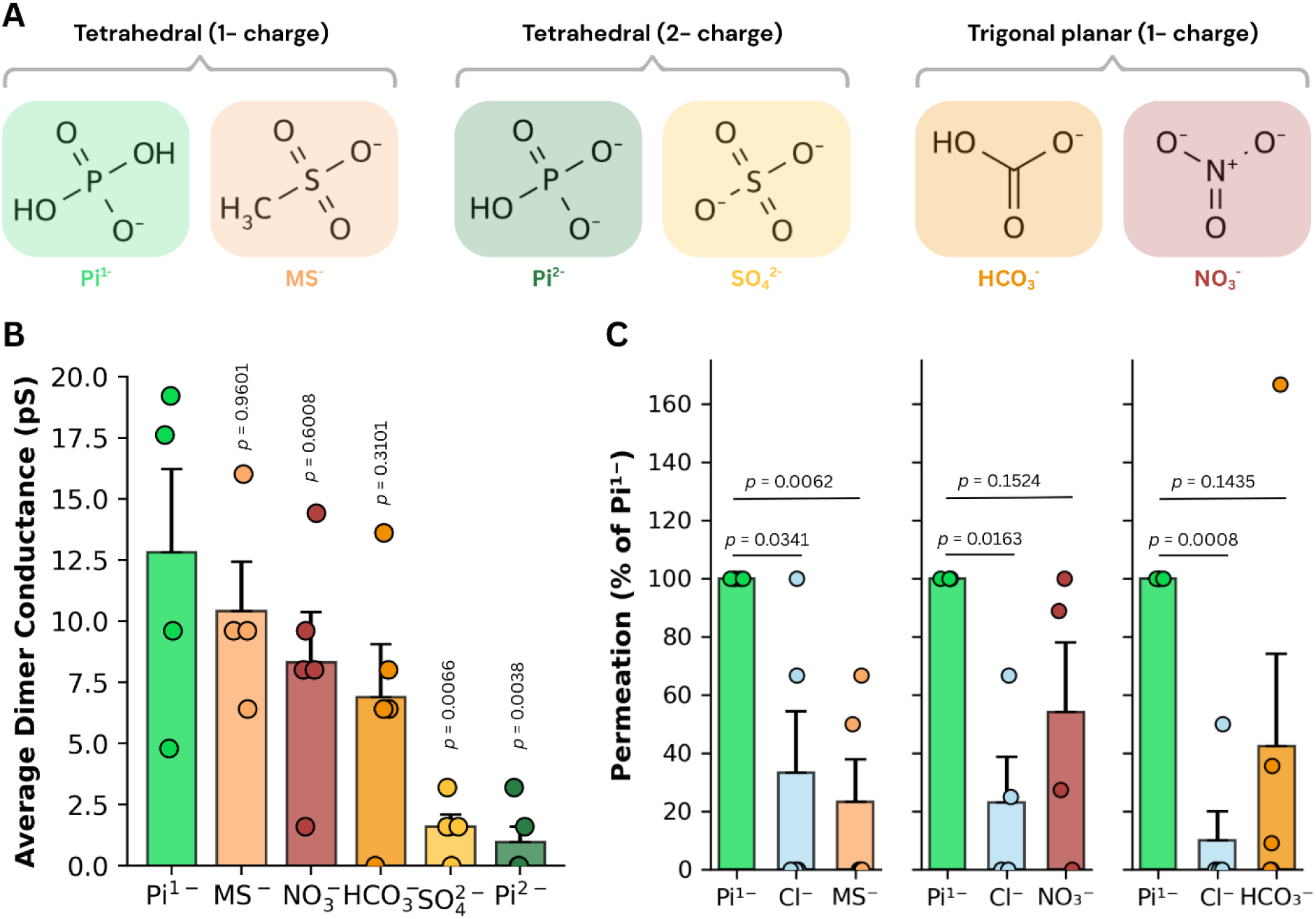
Phosphate competes with other monovalent anions for permeation through XPR1. **A)** Structures of tetrahedral monovalent anions Pi^1-^ and methanesulfonate (MS^-^), tetrahedral divalent anions Pi^2-^ and sulfate (SO_4_^2-^), and trigonal planar monovalent anions bicarbonate (HCO_3_^-^) and nitrate (NO_3_^-^). **B)** Average dimer conductance (pS) measured for each anion in simulations performed in the absence of phosphate. Conductance for Pi^1-^ and Pi^2-^ are shown as a reference. Bars represent the mean conductance across replicate simulations with individual replicate values overlaid. Error bars represent SEM. Statistical significance was determined using a one-way ANOVA with a post-hoc Tukey test, with *p* values for comparisons to Pi^1-^ shown above each bar. **C)** Relative permeation of competing anions in simulations containing Pi^1-^, Cl^-^, and the indicated competing anion. Bars show the percentage of Pi^1-^ and competing-anion permeation events during the final 1 μs of the simulations. Individual replicate values are overlaid. Error bars represent SEM. Statistical significance was assessed using paired *t*-tests comparing Cl^-^ and the competing anion with Pi^1-^ within each simulation system, and *p* values are shown above the corresponding comparisons.

We next measured permeation events for methanesulfonate, bicarbonate, and nitrate in the presence of Pi^1-^. In all three systems, Pi^1-^ had the greatest mean number of permeation events compared with both Cl^-^ and the competing anion. Pi^1-^ permeated significantly more frequently than Cl^-^ across all systems. However, the difference in permeation between Pi^1-^ and the competing anion was only statistically significant in the methanesulfonate system, while the number of permeation events observed for nitrate and bicarbonate did not differ significantly from Pi^1-^ (**Fig. 3C, Fig. S3A-C**). Analysis of RDFs for nitrate, bicarbonate, and methanesulfonate showed that these anions can each come into close contact with neighbouring ions of the same type, similar to Pi^1-^ (**Fig. S2**). Nevertheless, Pi^1-^ continued to permeate in all simulations, indicating that none of the competing anions prevented Pi conduction through XPR1.

## Discussion

Our previously proposed knock-kiss-kick mechanism^22^ shares several conceptual similarities with the ‘hard knock-on’ permeation mechanisms described for potassium channels, in which multiple ions simultaneously occupy the permeation pathway and direct electrostatic interactions between neighbouring ions facilitate conduction.^65–67^ However, important differences exist between these systems. In the hard knock-on mechanism, dehydrated K⁺ ions permeate through a carbonyl-lined selectivity filter within the channel, whereas Pi permeation through XPR1 occurs within a relatively more dynamic pathway lined by flexible side chains and involves transient stabilisation by positively charged residues together with Pi-Pi hydrogen-bonding interactions. Rather than direct electrostatic repulsion alone, Pi permeation appears to result from a combination of constriction widening, transient binding, and displacement by subsequent incoming Pi ions.^22^

Interestingly, Pi^2-^ was conducted much less frequently than Pi^1-^ (**Fig. 1A**). We previously showed Pi^2-^ preferentially accumulated around the exterior of XPR1, particularly on the intracellular side, and rarely progressed through the transport pathway.^22^ As many charged lysine and arginine residues are located on the intracellular side of the protein, this behaviour may reflect stronger electrostatic interactions between Pi^2-^ and these residues, preventing Pi^2-^ from being readily displaced into the transport pathway.

Although this observation suggests that XPR1 preferentially conducts the monovalent Pi species, it should be interpreted with some caution. Classical fixed-charge force fields have previously been reported to overestimate electrostatic interactions involving phosphate groups due to the absence of explicit electronic polarisation.^68,69^ Consequently, the tendency of HPO_4_^2-^ to associate with positively charged regions of XPR1 may be exaggerated in our simulations. Further, our classical molecular dynamics simulations do not explicitly model changes in protonation state. In our simulations, Pi is fixed as either H_2_PO_4_^-^ or HPO_4_^2-^. However, it is possible that changes in Pi protonation state occur during permeation through XPR1. Constant-pH molecular dynamics^70^ or hybrid QM/MM^71^ approaches could be used in future studies to investigate whether proton transfer contributes to phosphate conduction.

Similar to Pi^2-^, sulfate also exhibited negligible permeation in our simulations. This result is consistent with electrophysiology data from Lu et al. (2024), which showed that XPR1 negligibly conducts sulfate.^23^ As two chemically distinct divalent anions were both shown to be poorly conducted by XPR1 in our simulations, this suggests that the protein may be selective against divalent anions. Cl^-^ permeation was also observed in our simulations. In the absence of Pi, Cl^-^ is readily permeated through XPR1 and occupies the Pi-1, Pi-2, and Pi-3 sites within the transport pathway. Intriguingly, Lu et al. (2024) showed that Pi generated substantially larger currents than Cl^-^ using inside-out patch clamp electrophysiology with either NMDG-Pi or NMDG-Cl solutions in HEK293T cells transfected with XPR1.^23^ Although this appears inconsistent with our raw permeation counts, the conditions are not directly comparable as we do not have simulations in which Pi^1-^ is the only anion, as Cl^-^ ions were always included in our simulation systems. Crucially, once Pi enters the pathway, Cl^-^ permeation progressively declines (**Fig. 1C**). Free energy calculations show that this arises because phosphate binds more favourably at the Pi-1 site. Furthermore, while Cl^-^ remain hydrated and separated by at least 5 Å (**Fig. 2H**), neighbouring Pi ions interact directly at 2 Å (**Fig. 2J**), allowing incoming phosphate to displace resident ions while blocking Cl^-^ entry to the Pi-0 site. If this cooperative mechanism occurs *in vivo*, XPR1 should exhibit anomalous mole fraction behavior analogous to that observed in voltage-gated calcium channels^72^, in which increasing Pi concentration in a Cl^-^ background would initially reduce current by blocking Cl^-^ permeation before increasing Pi occupancy promotes Pi transport.

This behaviour observed in our simulations may help explain seemingly contrasting findings from recent electrophysiological studies. Wu et al. (2025) reported that increasing extracellular Cl^-^ concentration did not alter XPR1 currents when Pi was present, while Lu et al. (2024) showed that XPR1 can conduct Cl^-^ when Pi is absent.^23,36^ Our simulations suggest that once Pi occupies the transport pathway, it progressively outcompetes Cl^-^. Consequently, increasing Cl^-^ concentration alone may be insufficient to enhance XPR1 conductance when Pi is already occupying the pore.

Additionally, Lu et al. (2024) showed that nitrate could be conducted by XPR1 but permeated significantly less than Pi. As our simulations do not include the protein’s intracellular SPX domain, this domain may contribute to additional selectivity for other ions. Wu et al. (2025) also demonstrated that methanesulfonate can be conducted by XPR1 and suggested that XPR1 did not show selectivity between Pi and methanesulfonate.^36^ In our simulations, no significant difference in permeation was observed between Pi and methanesulfonate when either anion was simulated in the absence of the other. However, when both anions were simulated together, methanesulfonate permeated significantly less frequently than Pi. These findings suggest that, although methanesulfonate is readily conducted by XPR1, phosphate is preferentially conducted when the two substrates compete for occupancy of the transport pathway under the simulated conditions.

Interestingly, substrate promiscuity is not unique to the SPX-EXS family. The SPX-MFS protein VPT1 has also been reported to transport multiple inorganic anions, including phosphate, sulfate, nitrate and Cl^-^. Although VPT1 is predicted to operate through a classical MFS alternating-access mechanism^73,74^ rather than the channel-like mechanism proposed here for XPR1, these observations suggest that broad anion selectivity may be a more general feature of SPX-containing phosphate transport proteins.

Together, these observations suggest that, in the absence of the SPX domain, XPR1 does not achieve substrate selectivity through absolute exclusion of competing anions from the transport pathway. Instead, selectivity appears to arise from differential stabilisation of substrates upon entry into the transport pathway, and the impact of resident ions on incoming ones.

This mode of selectivity may be particularly well-suited to Pi transport. Unlike monatomic ions such as K⁺ or Na⁺, Pi is a large, polyatomic anion. Rather than a relatively rigid selectivity filter that excludes competing substrates, XPR1 has a fairly permissive permeation pathway that allows several monovalent anions to pass. The apparent ability of XPR1 to conduct multiple anions also raises the question of why XPR1 has not evolved to exclusively conduct Pi. Under physiological conditions, Pi, Cl^-^, and bicarbonate are all abundant intracellular ions^75,76^, whereas nitrate and methanesulfonate are less likely to be present in significant amounts. The level of selectivity for Pi over other anions that we observed in our simulations may represent a balance between efficient Pi export through XPR1 and the structural constraints of transporting polyatomic anions, rather than a requirement for strict substrate specificity within the XPR1 transport pathway.

However, as noted earlier, these simulations used only the transmembrane core and EXS domain, removing the intracellular SPX domain to reduce computational cost. Additional selectivity may therefore arise outside the simulated permeation pathway, particularly near the intracellular entrance where the SPX domain and associated regulatory interactions could influence substrate access, local electrostatics, or gating. This may be especially relevant for bicarbonate, which permeated at rates comparable to phosphate in the present simulations. Future simulations of full-length XPR1, ideally in complex with InsP₈ and any other regulatory partners of XPR1, will be needed to determine whether the SPX domain contributes to substrate discrimination.

Together, these findings demonstrate that substrate selectivity in XPR1 emerges from its knock–kiss–kick permeation mechanism. The unique ability of Pi^1-^ to form close, direct interactions with both the protein and neighbouring substrate ions drives efficient conduction while restricting competing anions. These findings provide a molecular basis for substrate discrimination in XPR1 and demonstrate how selectivity can emerge from the collective behaviour of multiple ions within a channel-like transport pathway.

## Acknowledgements

This research was undertaken with assistance from resources and services provided by the National Computational Infrastructure (NCI), supported by the Australian Government, under project g15 (B.C.), accessed through the national Merit Allocation Scheme (NCMAS) and the Australian National University Merit Allocation Scheme (ANUMAS). B.C. acknowledges funding from the Australian Research Council (DP250100893). C.W. was supported by an Australian Government Research Training Program (RTP) PhD Scholarship.

## References

1. Peacock, M. Phosphate Metabolism in Health and Disease. Calcif. Tissue Int. 108, 3–15 (2021).

2. Berndt, T. & Kumar, R. Phosphatonins and the regulation of phosphate homeostasis. Annu. Rev. Physiol. 69, 341–359 (2007).

3. Razzaque, M. S. Phosphate toxicity: new insights into an old problem. Clin. Sci. (Lond). 120, 91–97 (2011).

4. Forster, I. C., Hernando, N., Biber, J. & Murer, H. Phosphate transporters of the SLC20 and SLC34 families. Mol. Aspects Med. 34, 386–395 (2013).

5. Levi, M. et al. Mechanisms of phosphate transport. Nat. Rev. Nephrol. 15, 482–500 (2019).

6. Tsai, J. Y. et al. Structure of the sodium-dependent phosphate transporter reveals insights into human solute carrier SLC20. Sci. Adv. 6, (2020).

7. Giovannini, D., Touhami, J., Charnet, P., Sitbon, M. & Battini, J. L. Inorganic Phosphate Export by the Retrovirus Receptor XPR1 in Metazoans. Cell Rep. 3, 1866–1873 (2013).

8. Wege, S. & Poirier, Y. Expression of the mammalian Xenotropic Polytropic Virus Receptor 1 (XPR1) in tobacco leaves leads to phosphate export. FEBS Lett. 588, 482–489 (2014).

9. Jennings, M. L. Role of transporters in regulating mammalian intracellular inorganic phosphate. Front. Pharmacol. 14, (2023).

10. Legati, A. et al. Mutations in XPR1 cause primary familial brain calcification associated with altered phosphate export. Nat. Genet. 47, 579–581 (2015).

11. Anheim, M. et al. XPR1 mutations are a rare cause of primary familial brain calcification. J. Neurol. 263, 1559–1564 (2016).

12. Ansermet, C. et al. Renal fanconi syndrome and hypophosphatemic rickets in the absence of xenotropic and polytropic retroviral receptor in the nephron. Journal of the American Society of Nephrology 28, 1073–1078 (2017).

13. Yao, X. P. et al. Analysis of gene expression and functional characterization of XPR1: a pathogenic gene for primary familial brain calcification. Cell Tissue Res. 370, 267–273 (2017).

14. López-Sánchez, U. et al. Characterization of XPR1/SLC53A1 variants located outside of the SPX domain in patients with primary familial brain calcification. Scientific Reports *2019* 9:1 9, 6776- (2019).

15. Akasu-Nagayoshi, Y. et al. PHOSPHATE exporter XPR1/SLC53A1 is required for the tumorigenicity of epithelial ovarian cancer. Cancer Sci. 113, 2034–2043 (2022).

16. Bondeson, D. P. et al. Phosphate dysregulation via the XPR1-KIDINS220 protein complex is a therapeutic vulnerability in ovarian cancer. Nat. Cancer 3, 681–695 (2022).

17. Orimo, K. et al. A Japanese family with idiopathic basal ganglia calcification carrying a novel XPR1 variant. J. Neurol. Sci. 451, (2023).

18. Yan, R. et al. Human XPR1 structures reveal phosphate export mechanism. Nature *2024 633*:8031 633, 960–967 (2024).

19. He, Q. et al. Structural basis of phosphate export by human XPR1. Nature Communications *2025 16:1* 16, 683- (2025).

20. Wang, Y. et al. Structural basis of phosphate export by human XPR1. Cell Res. 35, 313– 316 (2025).

21. Chen, L., He, J., Wang, M. & She, J. Structure and function of human XPR1 in phosphate export. Nature Communications *2025* 16:1 **16**, 2983- (2025).

22. Wang, X. et al. KIDINS220 and InsP8 safeguard the stepwise regulation of phosphate exporter XPR1. Mol. Cell 85, 3209–3224.e8 (2025).

23. Lu, Y. et al. Structural basis for inositol pyrophosphate gating of the phosphate channel XPR1. Science *(1979)*. 386, (2024).

24. Zhang, W. et al. Structural insights into the mechanism of phosphate recognition and transport by XPR1. Nature Communications 16, 1–10 (2025).

25. Zhu, Q., Yaggi, M. F., Jork, N., Jessen, H. J. & Diver, M. M. Transport and InsP8 gating mechanisms of the human inorganic phosphate exporter XPR1. Nature Communications *2025 16:1* 16, 2770- (2025).

26. Zuo, P. et al. Synergistic activation of the human phosphate exporter XPR1 by KIDINS220 and inositol pyrophosphate. 16, 2879- (2025).

27. Wild, R. et al. Control of eukaryotic phosphate homeostasis by inositol polyphosphate sensor domains. Science 352, 986–990 (2016).

28. Li, X. et al. Control of XPR1-dependent cellular phosphate efflux by InsP8 is an exemplar for functionally-exclusive inositol pyrophosphate signaling. Proc. Natl. Acad. Sci. U. S. A. 117, 3568–3574 (2020).

29. Moritoh, Y. et al. The enzymatic activity of inositol hexakisphosphate kinase controls circulating phosphate in mammals. Nat. Commun. 12, (2021).

30. Wege, S. et al. The EXS Domain of PHO1 Participates in the Response of Shoots to Phosphate Deficiency via a Root-to-Shoot Signal. Plant Physiol. 170, 385–400 (2016).

31. López-Sánchez, U. et al. Interplay between primary familial brain calcification-associated SLC20A2 and XPR1 phosphate transporters requires inositol polyphosphates for control of cellular phosphate homeostasis. J. Biol. Chem. 295, 9366–9378 (2020).

32. Li, X. et al. Homeostatic coordination of cellular phosphate uptake and efflux requires an organelle-based receptor for the inositol pyrophosphate IP8. Cell Rep. 43, (2024).

33. Fang, S. et al. Structural mechanism underlying PHO1;H1-mediated phosphate transport in Arabidopsis. Nat. Plants 11, 309–320 (2025).

34. Xu, C. et al. A phosphate-sensing organelle regulates phosphate and tissue homeostasis. Nature 617, 798–806 (2023).

35. Yang, H. et al. Structural insights into the gating mechanism of the fission yeast phosphate exporter SpXpr1. Cell Discovery *2026 12:1* 12, 27- (2026).

36. Wu, H. et al. The identification of XPR1 as a voltage- and phosphate-activated phosphate-permeable ion channel. Nature Communications *2025 16:1* 16, 4519- (2025).

37. Lee, J. et al. CHARMM-GUI Input Generator for NAMD, GROMACS, AMBER, OpenMM, and CHARMM/OpenMM Simulations Using the CHARMM36 Additive Force Field. J. Chem. Theory Comput. 12, 405–413 (2016).

38. Jo, S., Kim, T., Iyer, V. G. & Im, W. CHARMM-GUI: A web-based graphical user interface for CHARMM. J. Comput. Chem. 29, 1859–1865 (2008).

39. Lomize, M. A., Pogozheva, I. D., Joo, H., Mosberg, H. I. & Lomize, A. L. OPM database and PPM web server: resources for positioning of proteins in membranes. Nucleic Acids Res. 40, (2012).

40. Emsley, P. & Cowtan, K. Coot: model-building tools for molecular graphics. Acta Crystallogr. D Biol. Crystallogr. 60, 2126–2132 (2004).

41. Liebschner, D. et al. Macromolecular structure determination using X-rays, neutrons and electrons: recent developments in Phenix. Acta Crystallogr. D Struct. Biol. 75, 861–877 (2019).

42. Abraham, M. J. et al. GROMACS: High performance molecular simulations through multi-level parallelism from laptops to supercomputers. SoftwareX 1–2, 19–25 (2015).

43. Huang, J. et al. CHARMM36m: An Improved Force Field for Folded and Intrinsically Disordered Proteins. Nat. Methods 14, 71 (2017).

44. Khan, H. M., MacKerell, A. D. & Reuter, N. Cation-π interactions between methylated ammonium groups and tryptophan in the CHARMM36 additive force field. J. Chem. Theory Comput. 15, 7 (2018).

45. Hess, B., Bekker, H., Berendsen, H. J. C. & Fraaije, J. G. E. M. LINCS: A Linear Constraint Solver for Molecular Simulations. J Comput Chem 18, 14631472 (1997).

46. Darden, T., York, D. & Pedersen, L. Particle mesh Ewald: An N⋅log(N) method for Ewald sums in large systems. J. Chem. Phys. 98, 10089–10092 (1993).

47. Berendsen, H. J. C., Postma, J. P. M., Van Gunsteren, W. F., Dinola, A. & Haak, J. R. Molecular dynamics with coupling to an external bath. J. Chem. Phys. 81, 3684–3690 (1984).

48. Bussi, G., Donadio, D. & Parrinello, M. Canonical sampling through velocity rescaling. Journal of Chemical Physics 126, 14101 (2007).

49. Parrinello, M. & Rahman, A. Polymorphic transitions in single crystals: A new molecular dynamics method. J. Appl. Phys. 52, 7182–7190 (1981).

50. Malde, A. K. et al. An Automated Force Field Topology Builder (ATB) and Repository: Version 1.0. J. Chem. Theory Comput. 7, 4026–4037 (2011).

51. Vanommeslaeghe, K. et al. CHARMM General Force Field (CGenFF): A force field for drug-like molecules compatible with the CHARMM all-atom additive biological force fields. J. Comput. Chem. 31, 671 (2010).

52. Shirts, M. R. & Chodera, J. D. Statistically optimal analysis of samples from multiple equilibrium states. Journal of Chemical Physics 129, 124105 (2008).

53. Thomas, M., Jayatilaka, D. & Corry, B. Mapping the Importance of Four Factors in Creating Monovalent Ion Selectivity in Biological Molecules. Biophys. J. 100, 60–69 (2011).

54. Bennett, C. H. Efficient Estimation of Free Energy Differences from Monte Carlo Data. J. Comput. Phys. 22, 245–268 (1976).

55. Miller, B. R. et al. MMPBSA.py: An Efficient Program for End-State Free Energy Calculations. J. Chem. Theory Comput. 8, 3314–3321 (2012).

56. Valdés-Tresanco, M. S., Valdés-Tresanco, M. E., Valiente, P. A. & Moreno, E. gmx_MMPBSA: A New Tool to Perform End-State Free Energy Calculations with GROMACS. J. Chem. Theory Comput. 17, 6281–6291 (2021).

57. Adamson, L. S. R. et al. Pore structure controls stability and molecular flux in engineered protein cages. Sci. Adv. 8, (2022).

58. Humphrey, W., Dalke, A. & Schulten, K. VMD: visual molecular dynamics. J. Mol. Graph. 14, 33–38 (1996).

59. Michaud-Agrawal, N., Denning, E. J., Woolf, T. B. & Beckstein, O. MDAnalysis: A toolkit for the analysis of molecular dynamics simulations. J. Comput. Chem. 32, 2319–2327 (2011).

60. Mckinney, W. Data Structures for Statistical Computing in Python. (2010).

61. Harris, C. R. et al. Array programming with NumPy. Nature *2020 585:* 7825 585, 357–362 (2020).

62. Hunter, J. Matplotlib: A 2D Graphics Environment. Comput. Sci. Eng. 9, (2007).

63. Waskom, M. L. seaborn: statistical data visualization. J. Open Source Softw. 6, 3021 (2021).

64. Virtanen, P. et al. SciPy 1.0: fundamental algorithms for scientific computing in Python. Nature Methods *2020 17:3* 17, 261–272 (2020).

65. Hodgkin, A. L. & Keynes, R. D. The potassium permeability of a giant nerve fibre. J. Physiol. 128, 61–88 (1955).

66. Doyle, D. A. et al. The structure of the potassium channel: molecular basis of K+ conduction and selectivity. Science 280, 69–77 (1998).

67. Köpfer, D. A. et al. Ion permeation in K^+^ channels occurs by direct Coulomb knock-on. Science 346, 352–355 (2014).

68. Puyo-Fourtine, J., Juillé, M., Hénin, J., Clavaguéra, C. & Duboué-Dijon, E. Consistent Picture of Phosphate–Divalent Cation Binding from Models with Implicit and Explicit Electronic Polarization. J. Phys. Chem. B 126, 4022–4034 (2022).

69. Korn, V. et al. Evaluating Force Matching as a Parametrization Strategy for the CHARMM36m Force Field Using Phosphorylation. J. Phys. Chem. B 130, 5469–5487 (2026).

70. Aho, N. et al. Scalable Constant pH Molecular Dynamics in GROMACS. J. Chem. Theory Comput. 18, 6148–6160 (2022).

71. Morozov, D. et al. Seamless QM/MM simulations via a GROMACS–CP2K interface. https://doi.org/10.26434/CHEMRXIV.15005148/V1 (2026) doi:10.26434/CHEMRXIV.15005148/V1.

72. Gillespie, D. & Boda, D. The Anomalous Mole Fraction Effect in Calcium Channels: A Measure of Preferential Selectivity. Biophys. J. 95, 2658 (2008).

73. Liu, J. et al. A vacuolar phosphate transporter essential for phosphate homeostasis in Arabidopsis. Proc. Natl. Acad. Sci. U. S. A. 112, E6571–E6578 (2015).

74. Wang, C. et al. Rice SPX-Major Facility Superfamily3, a Vacuolar Phosphate Efflux Transporter, Is Involved in Maintaining Phosphate Homeostasis in Rice. Plant Physiol. 169, pp.01005.2015 (2015).

75. Raut, S. K. et al. Chloride ions in health and disease. Biosci. Rep. 44, BSR20240029 (2024).

76. Khuri, R. N., Agulian, S. K. & Bogharian, K. K. Intracellular bicarbonate of skeletal muscle under different metabolic states. Am. J. Physiol. 230, 228–232 (1976).

