## Supplementary Information for "Phosphate selectivity in XPR1 is driven by direct multi-ion competition"

### Supplementary Movies

**Movie S1. Replacement of $Cl^-$ by Pi within the transport pathway.** $Cl^-$ ions conduct at the start of iO-eO simulations before being replaced within the cavity by Pi, leading to increasingly fewer $Cl^-$ permeation events over time.

### Supplementary Material

**Table S1. Data from free energy perturbation calculations.** Forwards calculations refer to simulations where $Cl^-$ was transformed into Pi, while reverse transformations refer to simulations where Pi was transformed into $Cl^-$.

| System | Direction | ΔG (kJ/mol) | | | | |
|---|---|---|---|---|---|---|
| | | Rep 1 | Rep 2 | Rep 3 | Mean | SEM |
| Pi-0 | Forwards | -293.01 | -292.10 | -292.53 | -292.55 | 0.26 |
| Pi-0 | Reverse | 291.32 | 293.40 | 290.53 | 291.75 | 0.86 |
| Pi-1 | Forwards | -303.77 | -299.80 | -301.02 | -301.53 | 1.17 |
| Pi-1 | Reverse | 301.02 | 301.49 | 297.28 | 299.93 | 1.33 |
| Pi-0 (with Pi at Pi-1) | Forwards | -300.87 | -300.38 | -301.92 | -301.06 | 0.45 |
| Pi-0 (with Pi at Pi-1) | Reverse | 301.68 | 300.95 | 299.83 | 300.82 | 0.54 |
| Bulk | Forwards | -294.40 | -294.48 | -294.30 | -294.39 | 0.05 |
| Bulk | Reverse | 294.53 | 294.46 | 294.52 | 294.50 | 0.02 |

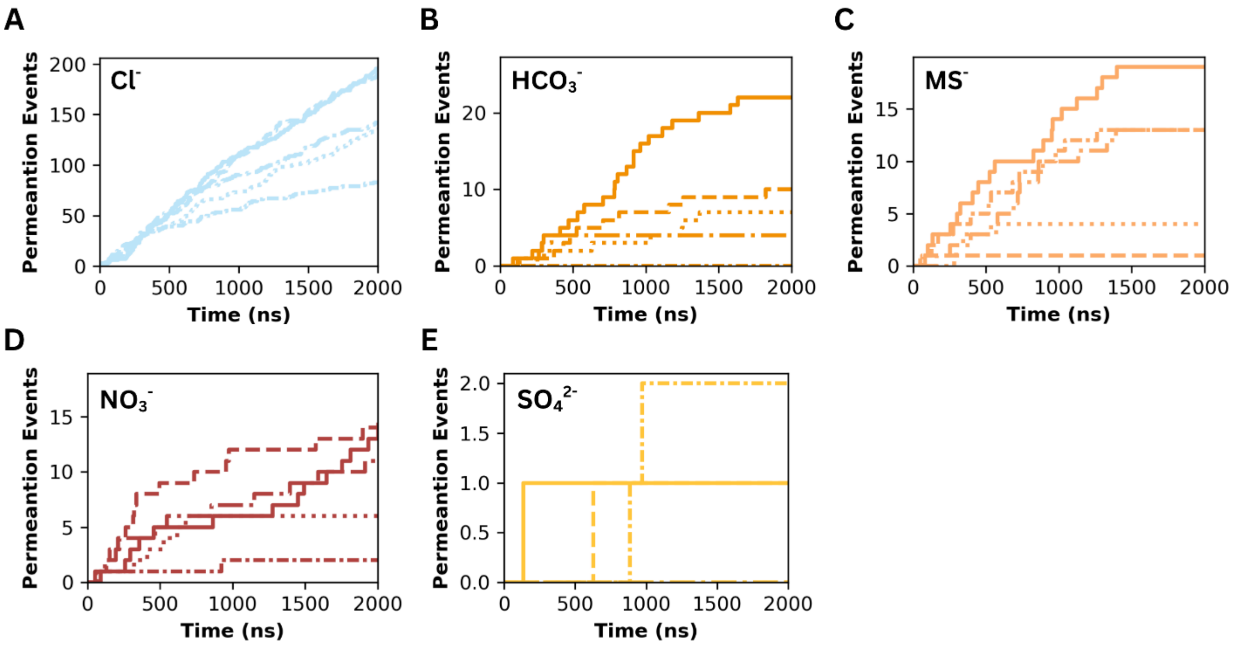


**Figure S1. Permeation events over time for anions in the absence of Pi.** Cumulative permeation events for **A)** $Cl^-$, **B)** $HCO_3^-$, **C)** $MS^-$, **D)** $NO_3^-$, **E)** $SO_4^{2-}$, recorded from MD simulations of the iO-eO structure in the absence of $Pi^{1-}$ at –400 mV. Each line represents one replicate.

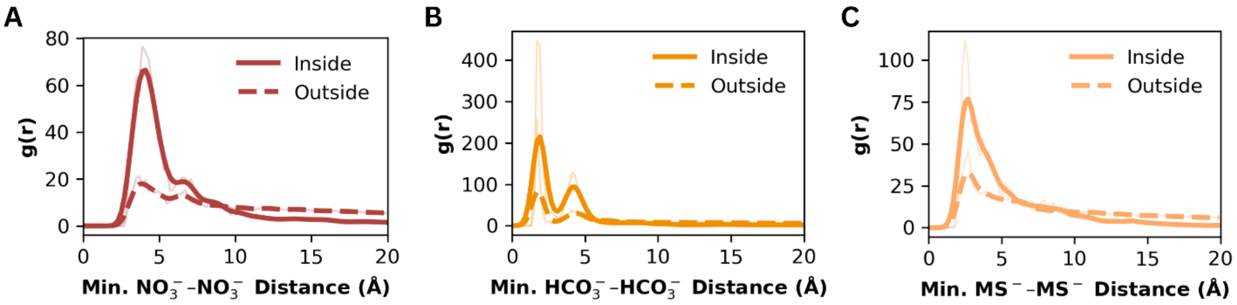


**Figure S2. Radial distribution functions for competing anions.** Radial distribution functions showing the distributions of minimum **A)** $NO_3^-$–$NO_3^-$, **B)** $HCO_3^-$–$HCO_3^-$, and **C)** $MS^-$–$MS^-$ distances for ions inside and outside the transport pathway. Traces for outside of the pathway in **A-C** were multiplied by 5 to be more visible within the panel.

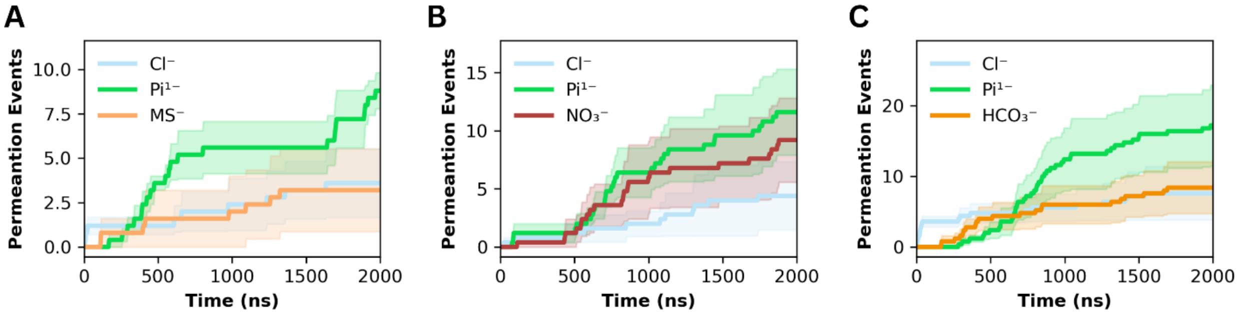


**Figure S3. Permeation events over time for anions in the presence of Pi.** Traces for the cumulative number of permeation events for $Cl^-$, $Pi^{1-}$, and the competing anion over 2 µs of simulation, where the anion was **A)** methanesulfonate ($MS^-$), **B)** nitrate ($NO_3^-$), or **C)** bicarbonate ($HCO_3^-$). Simulations were run using the iO-eO structure at -400 mV. For **A-C**, the traces represent the mean number of events across replicates. Shaded regions = SEM.

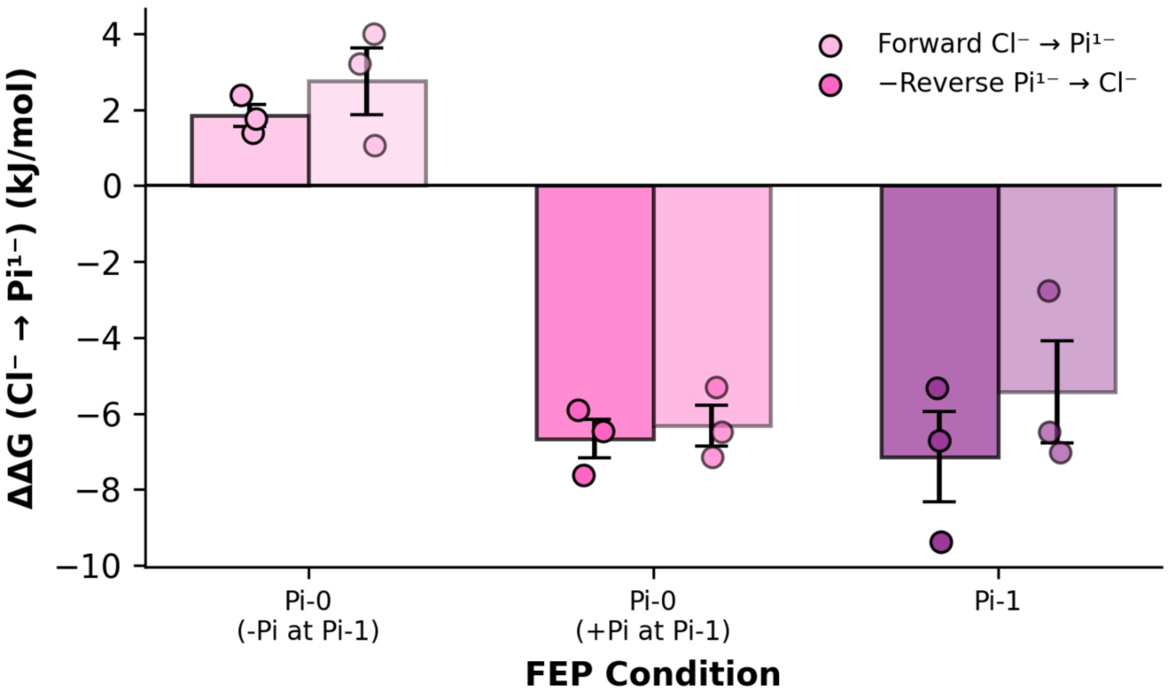


**Figure S4. Convergence of free energy perturbation simulations.** Comparison of ΔΔG values obtained from forward ($Cl^- \rightarrow H_2PO_4^-$) and reverse ($H_2PO_4^- \rightarrow Cl^-$) free energy perturbation calculations at the Pi-0 and Pi-1 binding sites. Calculations at Pi-0 were performed both in the absence and presence of $H_2PO_4^-$ at the Pi-1 site. Reverse ΔΔG values were sign-inverted to allow direct comparison with the forward calculations. Bars represent the mean of three independent replicates, with individual replicate values overlaid as scatter points. Error bars represent the standard error of the mean (SEM).

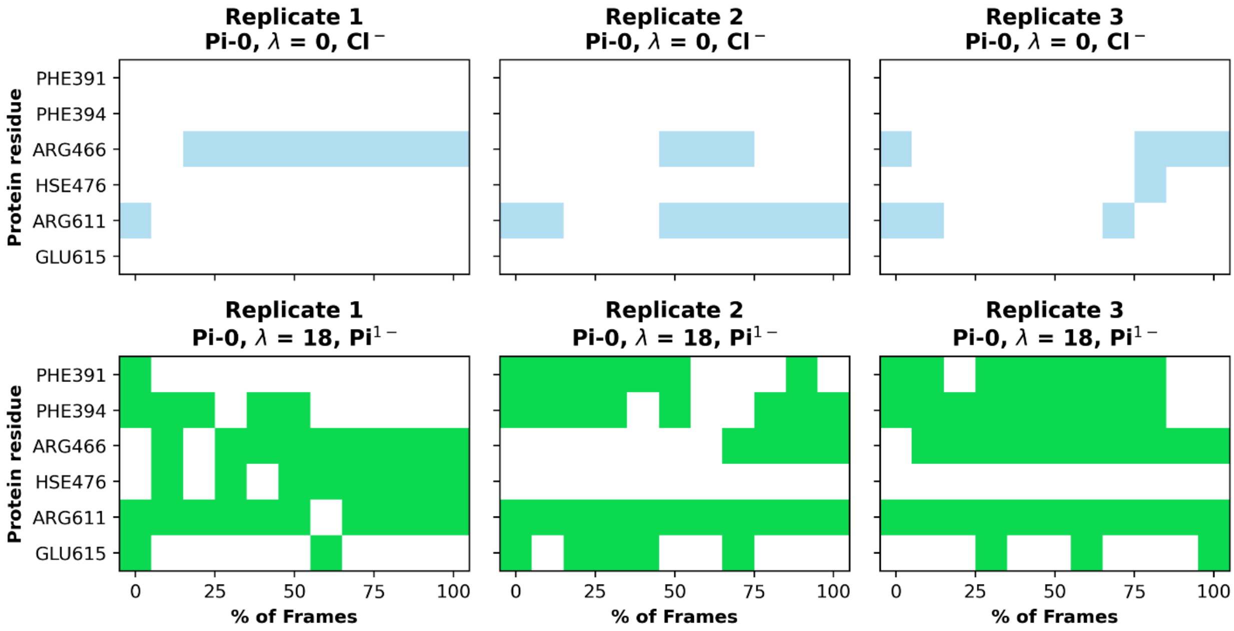


**Figure S5. Residue contacts over time from FEP at the Pi-0 site.** Residue contacts formed between the $Cl^-$ (window 0, λ = 0) and $Pi^{1-}$ (window 18, λ=1) from each replicate are shown over time. Each λ window for each replicate was simulated for 10 ns.

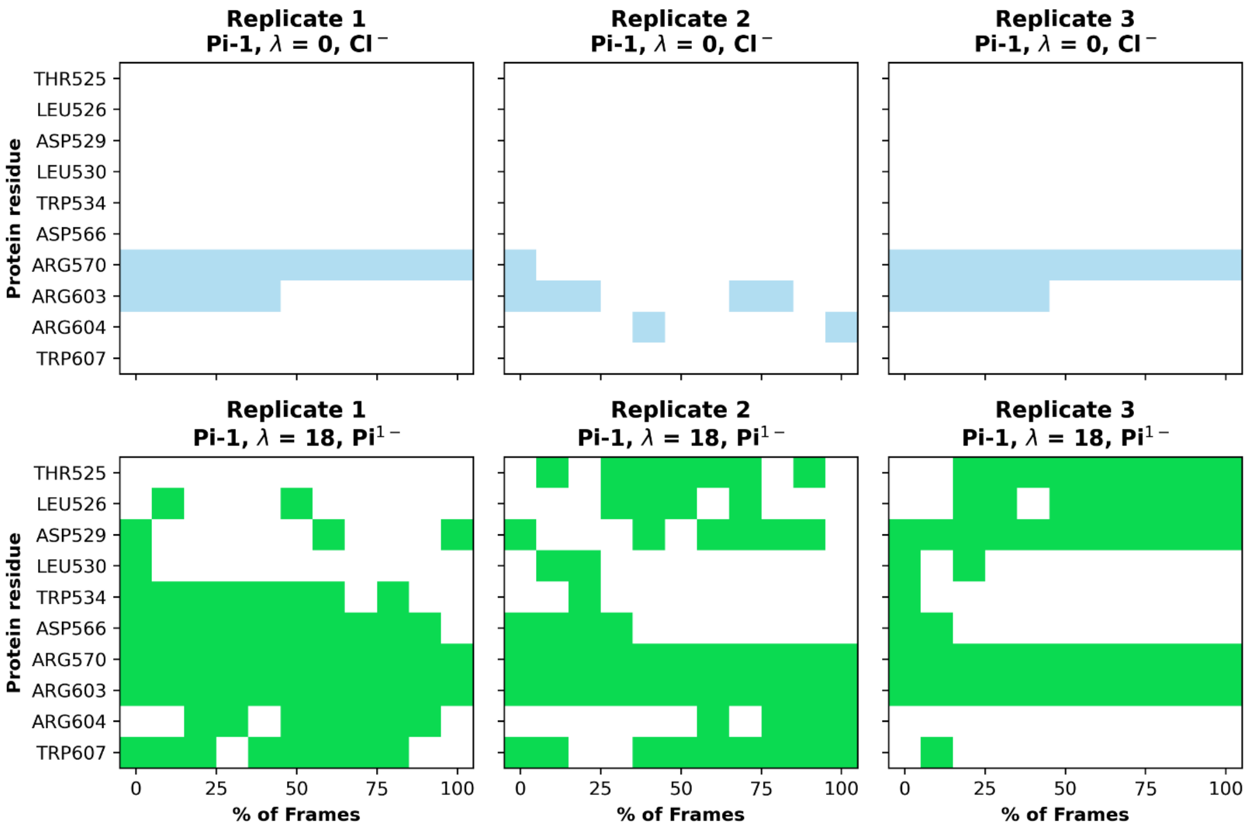


**Figure S6. Residue contacts over time from FEP at the Pi-1 site.** Residue contacts formed between the $Cl^{-}$ (window 0, λ = 0) and $Pi^{1-}$ (window 18, λ=1) from each replicate are shown over time. Each λ window for each replicate was simulated for 10 ns.

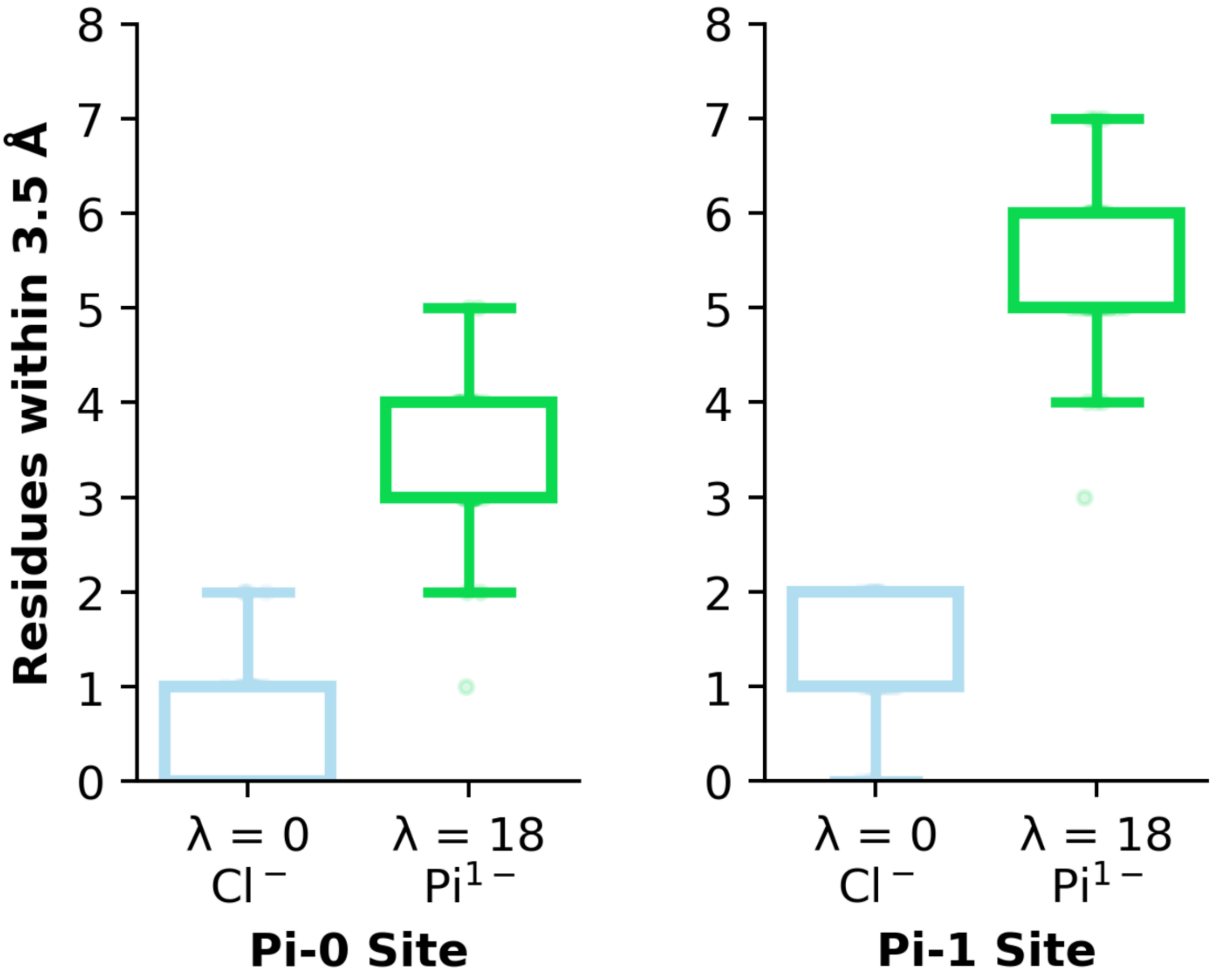


**Figure S7. Number of residue interactions for $Pi^{1-}$ and $Cl^{-}$ at the Pi-0 and Pi-1 sites.** Boxplots for the number of contacts formed in FEP simulations at the endpoints of λ = 0 (window 0) and λ = 1 (window 18) at the Pi-0 and Pi-1 sites. Residues within 3.5 Å of either the $Cl^{-}$ atom at λ = 0 (window 0) or the $Pi^{1-}$ atoms at λ = 1 (window 18) were measured across each frame of each replicate's trajectories. The residues forming the interactions are shown in Fig. S5 for the Pi-0 site and Fig. S6 for the Pi-1 site.

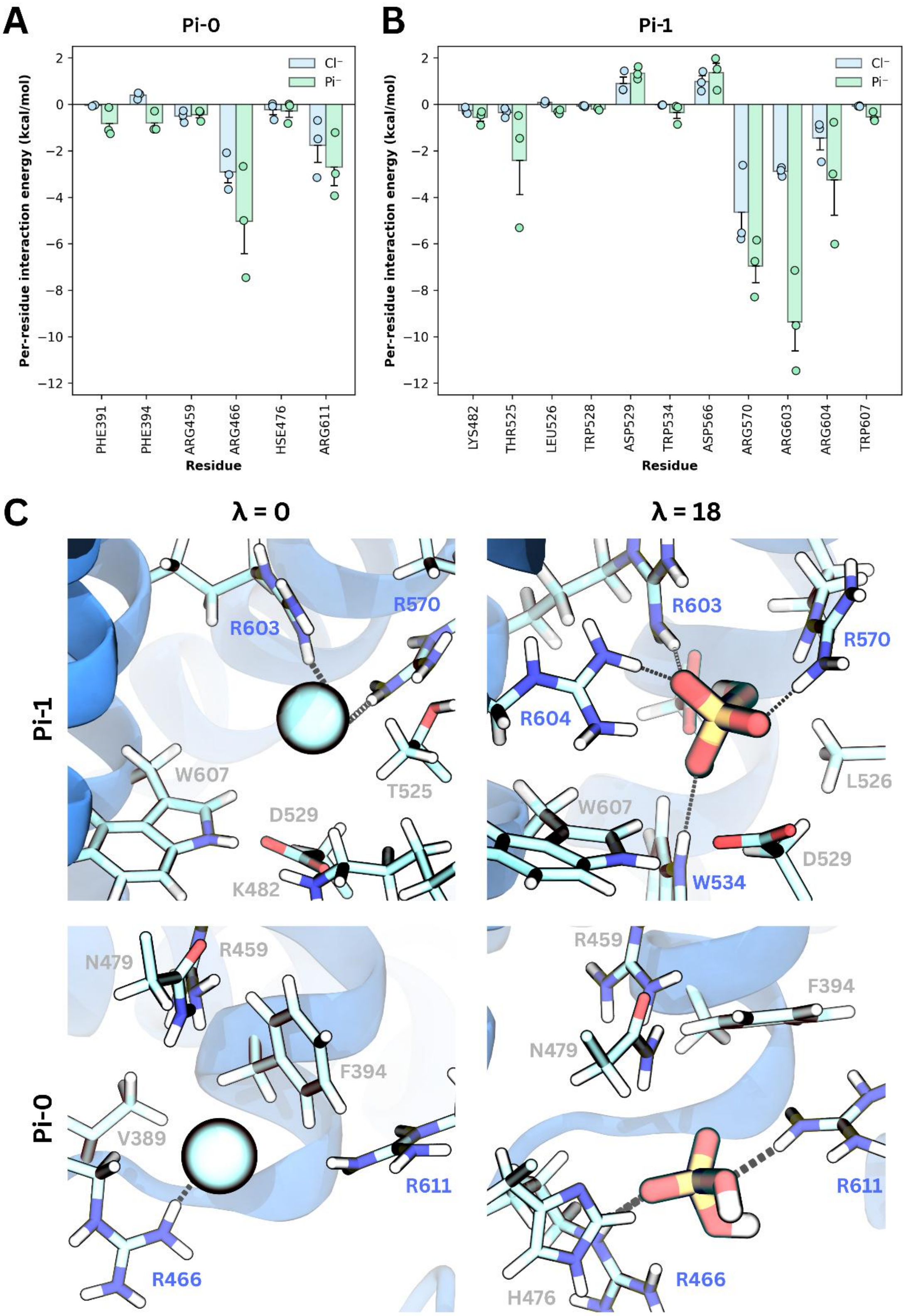


**Figure S8. Interactions formed by $Pi^{1-}$ and $Cl^{-}$ at the Pi-0 and Pi-1 sites. A–B)** Per-residue interaction energies (kcal/mol) calculated using MM/GBSA for the **A)** Pi-0 and **B)** Pi-1 binding sites. Interaction energies for $Cl^{-}$ are shown in blue, and those for $Pi^{1-}$ are shown in green. Bars represent the mean interaction energy across three independent replicates, with individual replicate values overlaid as points. Error bars represent SEM. $Pi^{1-}$ interaction energies were calculated from the λ = 1 (window 18) window of the forward FEP simulations, while $Cl^{-}$ interaction energies were calculated from the λ = 0 window of the forward FEP simulations. **C)** Representative snapshots from the $\lambda = 0$ and $\lambda = 1$ (window 18) windows at the Pi-0 and Pi-1 sites. Residues within 3.5 Å of the anion are shown. Residues positioned to form hydrogen bonds with $Cl^-$ or $Pi^{1-}$ in the representative snapshots are highlighted in blue. Hydrogen bonds predicted by VMD are shown as black dashed lines.
